# Accelerated Nitrogen Cycling on Seagrass Leaves in a High-CO_2_ World

**DOI:** 10.1101/2023.05.19.541481

**Authors:** Johanna Berlinghof, Luis M. Montilla, Friederike Peiffer, Grazia M. Quero, Ugo Marzocchi, Travis B. Meador, Francesca Margiotta, Maria Abagnale, Christian Wild, Ulisse Cardini

## Abstract

Seagrass meadows form highly productive and diverse ecosystems in coastal areas worldwide, where they are increasingly exposed to ocean acidification (OA). Efficient nitrogen (N) cycling and uptake are essential to maintain plant productivity, but the effects of OA on N transformations in these systems are poorly understood. Here we show that complete N cycling occurs on leaves of the Mediterranean seagrass *Posidonia oceanica*, with OA affecting both N gain and loss while the prokaryotic community structure remains largely unaffected. Daily leaf-associated N_2_ fixation contributed to 35% of the plant’s N demand under ambient pH, whereas it contributed to 45% under OA. Nitrification potential was only detected under OA, and N-loss via N_2_ production increased, although the balance remained decisively in favor of enhanced N gain. Our work highlights the role of the N-cycling microbiome in seagrass adaptation to OA, with key N transformations accelerating towards increased N gain.

## Introduction

Seagrass meadows are highly productive ecosystems worldwide, often occurring in nutrient-limited coastal areas^1^. They are among the most ecologically and economically valuable ecosystems on Earth^2^. Providing habitat, breeding grounds, and food for a wide range of organisms, they are considered ‘hotspots’ for biodiversity^3^. They also play an important role in sequestering large amounts of carbon, comparable to terrestrial forests^4^. In particular, the Mediterranean seagrass *Posidonia oceanica* can contribute to climate change mitigation through its effective CO_2_ uptake and large sequestration capacity^5^ and may even act as a buffer against ocean acidification (OA) by temporarily raising the seawater pH through its daylight photosynthesis^6^. This is relevant since the Mediterranean Sea has a higher capacity to absorb anthropogenic CO_2_ than other oceans due to its particular CO_2_ chemistry and active overturning circulation^7^.

Generally, marine plants are expected to benefit from increased CO_2_ concentrations as their photosynthetic rates are undersaturated at current CO_2_ levels^8^. However, OA has multifaceted effects on *P. oceanica*. Photosynthetic performance of *P. oceanica* seedlings and net leaf productivity increase under high pCO_2_^9–11^, while OA has little effect on the net community production of *P. oceanica* but results in increased shoot density and shorter leaf length due to increased herbivory^11–13^. Calcareous epiphytes such as encrusting red algae, bryozoans, foraminifers, and spirorbids decline or even disappear under OA, while non-calcareous invertebrates such as hydrozoans and tunicates benefit^11,14–16^.

Much less attention has been paid to the effects of OA on the biogeochemical cycling of elements other than carbon, such as nitrogen (N). Nitrogen is an essential nutrient for all living organisms and can be a limiting factor for primary production in marine seagrasses^17^, with its availability depending on diverse N transformation processes that are performed by a complex network of metabolically diverse microorganisms^18^. Seawater pH affects N speciation and concentration, which in turn affects metabolic processes and N transformations^19,20^. Dinitrogen (N_2_) fixation by N_2_-fixing bacteria and archaea (i.e., diazotrophs) has often been found to increase under OA^20,21^. The reason is not always clear, but in phototrophs, it may involve more energy being redirected to the demanding N_2_ fixation process owing to the down-regulation of carbon-concentrating mechanisms^20,22,23^. Autotrophic microbial nitrification can be highly sensitive to pH, and nitrification in the open ocean has been found to be significantly reduced by OA^24^. Dissimilatory nitrate reduction processes (e.g., denitrification or anaerobic ammonium oxidation -anammox), which are modular and involve many different bacterial groups often found in low-pH environments, are thought to be less affected by OA, with rates showing contrasting results at low seawater pH^20^.

Many N-cycling microorganisms can be found in close association with *P. oceanica*, together forming a holobiont^25,26^. N_2_ fixation by associated diazotrophic microorganisms can be crucial in providing the N required for seagrass photosynthesis and growth when its availability is limited^27,28^. Diazotrophic bacteria have been detected in the rhizosphere of *P. oceanica*^29^ with high rates of root-associated N_2_ fixation reported^30^. Analogous to many land plants that associate with diazotrophs, a recent study shows that *P. oceanica* lives in symbiosis with an N_2_-fixing γ-proteobacterium in its roots, providing N in exchange for sugars, that can fully sustain plant biomass production during its primary growth season^27^. Apart from this root-symbiosis, N_2_ fixation has been shown to occur associated with all parts of *P. oceanica*, both above and below ground^31^.

Overall, although rhizosphere N cycling has been the focus of extensive research, precise quantification of N transformations on seagrass leaves, as well as an evaluation of the effects of OA, are still lacking. Besides N_2_ fixation, we hypothesize that seagrass leaves could also be suitable sites for nitrification. For example, Ling et al.^32^ found a diverse community of ammonia-oxidizing archaea (AOA) and bacteria (AOB) associated with different parts of the seagrass *Thalassia hemprichii*, including leaf tissues. Moreover, anoxic parts within μm to mm-thick biofilms on the leaf surface could provide potential microhabitats for N loss pathways, such as denitrification^33,34^ or anammox performed by groups such as Planctomycetes, which were found to dominate the microbiome of *P. oceanica* leaves at some locations^35^.

Here, we investigate the effects of long-term natural OA on the epiphytic prokaryotic community of *P. oceanica* leaves and quantify rates of the key N cycling processes by the plant phyllosphere. We test the effects of pH and the presence/absence of epiphytes in multifactorial laboratory incubations, using N stable isotope tracers to quantify N_2_ fixation, nitrification potential, and anammox and denitrification potential, and net nutrient fluxes to quantify assimilatory processes by leaves and epiphytes. We complement these analyses with 16s rRNA gene amplicon sequencing to explore the diversity of the phyllosphere microbial community and the potential players involved in N transformation processes on seagrass leaves.

## Results and discussion

### Complete microbial N cycling occurs in the *P. oceanica* phyllosphere

Incubation experiments with ^15^N stable isotope labeling reveal that all key microbial N cycling processes occurred in the phyllosphere of *P. oceanica*, with epiphytes contributing to a net N gain in all conditions by the holobiont. To quantify rates of N_2_ fixation by the phyllosphere diazotrophic community, we incubated leaf sections with and without epiphytes in ^15^N_2_-enriched seawater. We detected clear epiphytic ^15^N_2_ incorporation in the light incubations, ranging from 0.12 ± 0.05 nmol cm^-2^ h^-1^ (mean ± SE) at the ambient site to 0.62 ± 0.15 nmol N cm^-2^ h^-1^ at the vent site (Fig. 1a). ^15^N_2_ incorporation was 370% higher at the vent site (F_1,12_ = 7.20, p = 0.020, R^2^ = 0.68) and in the same order of magnitude as N_2_ fixation rates measured *in situ* in minimally disturbed *P. oceanica* meadows^36^. Corresponding to dry weight-based rates of up to 131.08 nmol N g DW^-1^ h^-1^, these rates are also comparable to N_2_ fixation rates measured by root symbionts of *P. oceanica*^27,30^. Conversely, we observed significant ^15^N_2_ incorporation in only one of four replicates in the dark. We did not observe a significant transfer of fixed N to the *P. oceanica* plant tissues in the limited time frame of the experiment, neither in the light nor in the dark (Suppl. Figs. 2,3).

**Fig. 1.**
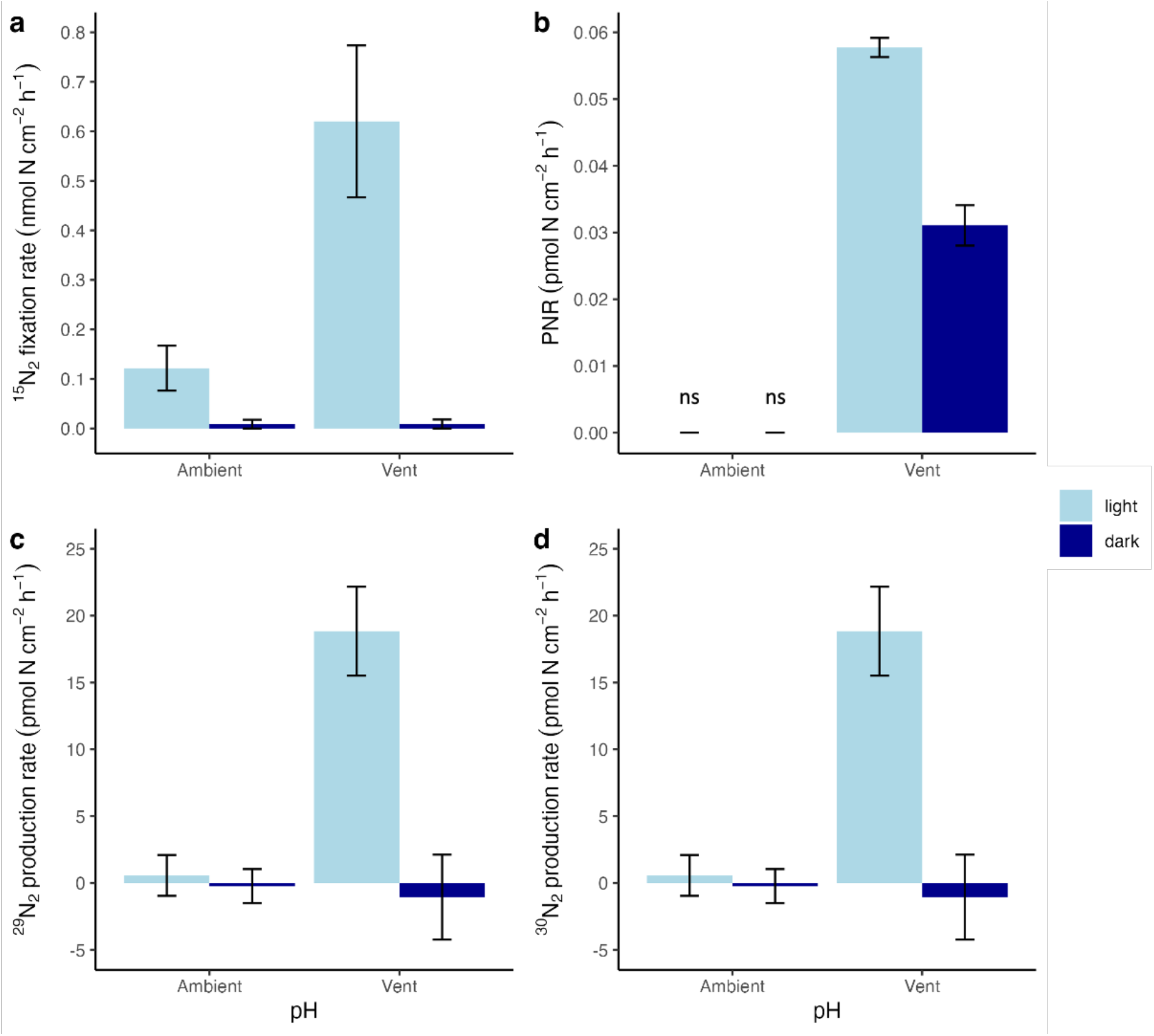
Nitrogen transformations in light and dark incubations from the ambient and vent site. Epiphytic ^15^N_2_ fixation rates (**a**; n = 4), potential nitrification rates (PNR) in incubations with leaf sections with epiphytes (**b**; n = 4), ^29^N_2_ (**c**) and ^30^N_2_ production rate (**d**) in incubations with leaf sections with epiphytes (n = 4). Error bars indicate mean ± SE, ns indicates enrichment was not significant.

Fixed N in the form of NH_4_^+^ can be recycled and converted into NO_3_^-^ by nitrification. We explored the potential of the phyllosphere microbiome to nitrify in ^15^N-NH_4_^+^ incubation experiments. While there was a strong variability among samples (Suppl. Fig. 4), we found significant (>2.5 x SD), but only marginal potential nitrification rates (PNR) at the vent site when epiphytes were present (Fig. 1b), ranging from 0.031 ± 0.003 pmol N cm^-2^ h^-1^ in the dark to 0.058 ± 0.0013 pmol N cm^-2^ h^-1^ in the light. PNR was 186% higher in the light (F_1,4_ = 63.48, p = 0.001, R^2^ = 0.99). In contrast, we found no significant PNR in incubations with epiphytes from the ambient site, neither in the light nor in the dark. The plant can compete with nitrifiers for N, as NH_4_^+^ is typically readily taken up by *P. oceanica*^37^, making the leaf phyllosphere a challenging environment for nitrifying prokaryotes. Our measurements of PNR in *P. oceanica* leaves are of relevance, as it indicates that a community of nitrifiers exists that can compete with the plant for NH_4_^+^ uptake. However, with PNR of up to 0.058 ± 0.0013 pmol N cm^-2^ h^-1^, their net contribution to NH_4_^+^ or NO_2_^-^ oxidation contributes only marginally to the N budget of the *P. oceanica* phyllosphere.

Previous studies suggested that anoxic parts within thick biofilms on the surface of seagrasses could be suitable microhabitats for N-loss pathways, such as denitrification and anammox^33,34^. Using incubation experiments of leaf sections amended with ^15^N-NO_3_^-^, we report ^29^N_2_ production rates ranging from 2.29 ± 0.53 pmol N cm^-2^ h^-1^ at the ambient site in the dark to 7.00 ± 2.07 pmol N cm^-2^ h^-1^ at the vent site in the light (Fig. 1c). ^29^N_2_ production was 234% higher at the vent site (F_1,13_ = 10.82, p = 0.006, R^2^ = 0.39), while the light/dark treatment had no effect. A significant production rate of ^30^N_2_ was only detected at the vent site in the light with 18.84 ± 3.33 pmol N cm^-2^ h^-1^ (Fig. 1d). Based on these results, we calculated daily budgets of total N-N_2_ loss (sum of ^29^N_2_ and ^30^N_2_ production, Fig. 4) of up to 4.01 ± 0.74 μmol N m^-2^ d^-1^ (or 0.401 ± 0.074 nmol N cm^-2^ d^-1^) at the vent site. These rates are significant, and comparable to N loss rates reported from seagrass sediments by Salk et al.^38^, who measured denitrification rates of 0.10 nmol N cm^-2^ d^-1^ and anammox rates of 0.43 nmol N cm^-2^ d^-1^. The presence of Planctomycetes and detectable rates of ^29^N_2_ in ^15^N-NO_3_^-^ amended incubations suggest that anammox may play an important role as an N loss pathway on seagrass leaves.

*P. oceanica* can assimilate fixed N as NH_4_^+^ or NO_3_^-37^ but shows a higher affinity for NH_4_^+39^. While NH_4_^+^ uptake rates were unaffected by the presence or absence of epiphytes (Suppl. Fig. 5 a, b), NO_3_^-^ consumption rates (Suppl. Fig. 5 c, d) were increased by 40 - 77 % in the presence of epiphytes. This is probably due to active NO_3_^-^ uptake because NO_3_^-^ loss via denitrification or anammox and nitrification activity was three orders of magnitude lower. Conversely to NH_4_^+^ uptake, NO_3_^-^ uptake rates were affected by the presence of epiphytes, suggesting that epiphytes may preferentially use this form of N as a strategy to avoid competition for N with the plant, combining active NO_3_^-^ uptake and N_2_ fixation.

### Distinct microbial communities contribute to seagrass phyllosphere N cycling

The 16s rRNA gene amplicon sequencing of the phyllosphere-associated microbiome revealed a diverse microbial community differing (see Suppl. Table 2) between the water column and seagrass leaf community and including many members potentially involved in N transformation processes on *P. oceanica* leaves.

The leaves were dominated by the phylum *Proteobacteria* with the classes *Alphaproteobacteria* (20-22%) and *Gammaproteobacteria* (9-15%) across both pH sites (Fig. 2). Among the predominant orders were *Rhodobacterales* (9%), which are commonly found as first colonizers on marine surfaces and seagrasses, probably due to their ability to be opportunistic and persist in rapidly changing environments ^40–42^. *Rhodobacterales* also include (putative) N_2_ fixers in both terrestrial^43^ and marine^44,45^ environments. We found *Rhizobiales* accounting for 5% of the total leaf community, a taxonomic order that includes a diversity of N_2_-fixing microbes that form symbiotic relationships with terrestrial plants ^46^ and known for promoting plant health and growth^47^.

*Cyanobacteria* accounted for 2-14% of the total leaf community (Fig. 2). Especially the orders *Phormidesmiales* and *Cyanobacteriales* had a large effect in the differential abundance analysis (Fig. 3). Higher N_2_ fixation rates under light conditions suggest a diazotrophic community dominated by species that can cope with O_2_ production from daytime photosynthesis, which would otherwise irreversibly inhibit the enzyme nitrogenase. Among the genera that can sustain N_2_ fixation in the light^48,49^, the leaves from both pH regimes comprised sequences for *Schizothrix* (0.22% on leaves vs. 0.01% in the water column) and *Trichodesmium* (up to 0.5% on leaves vs. 0.002% in the water column).

**Fig. 2.**
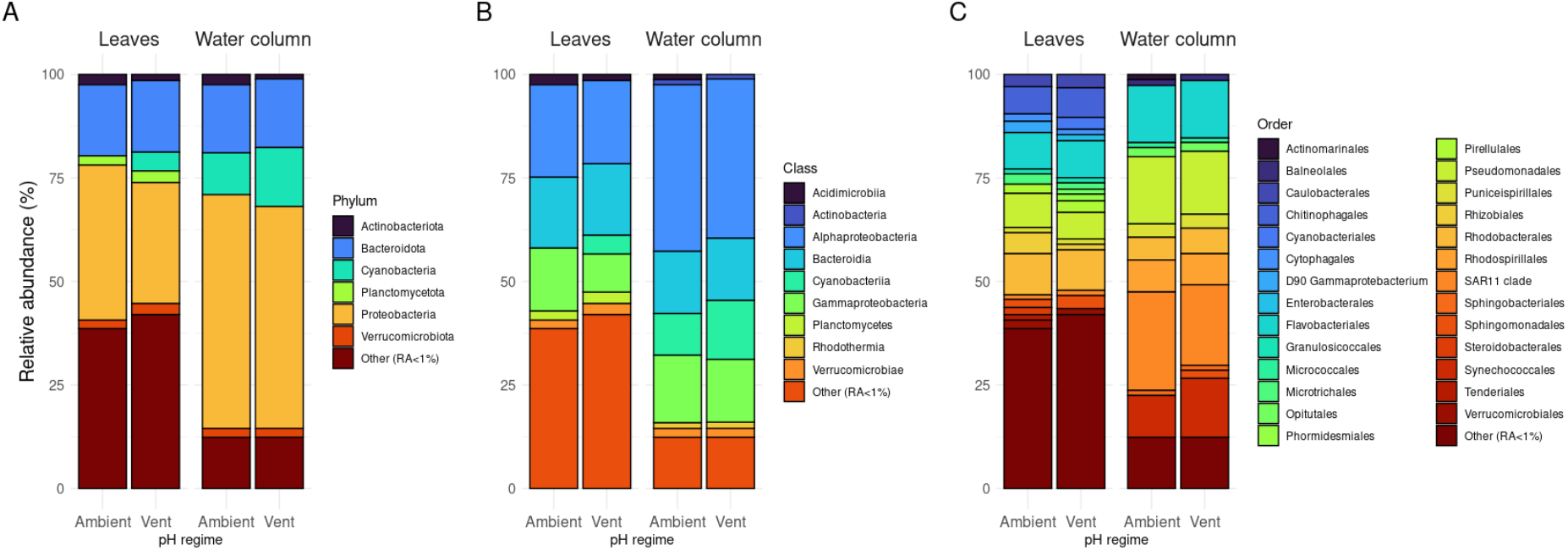
Average relative abundance of prokaryotic phyla (A), classes (B), and families (C) on leaves and water column samples from both pH regimes (n=3).

**Fig. 3.**
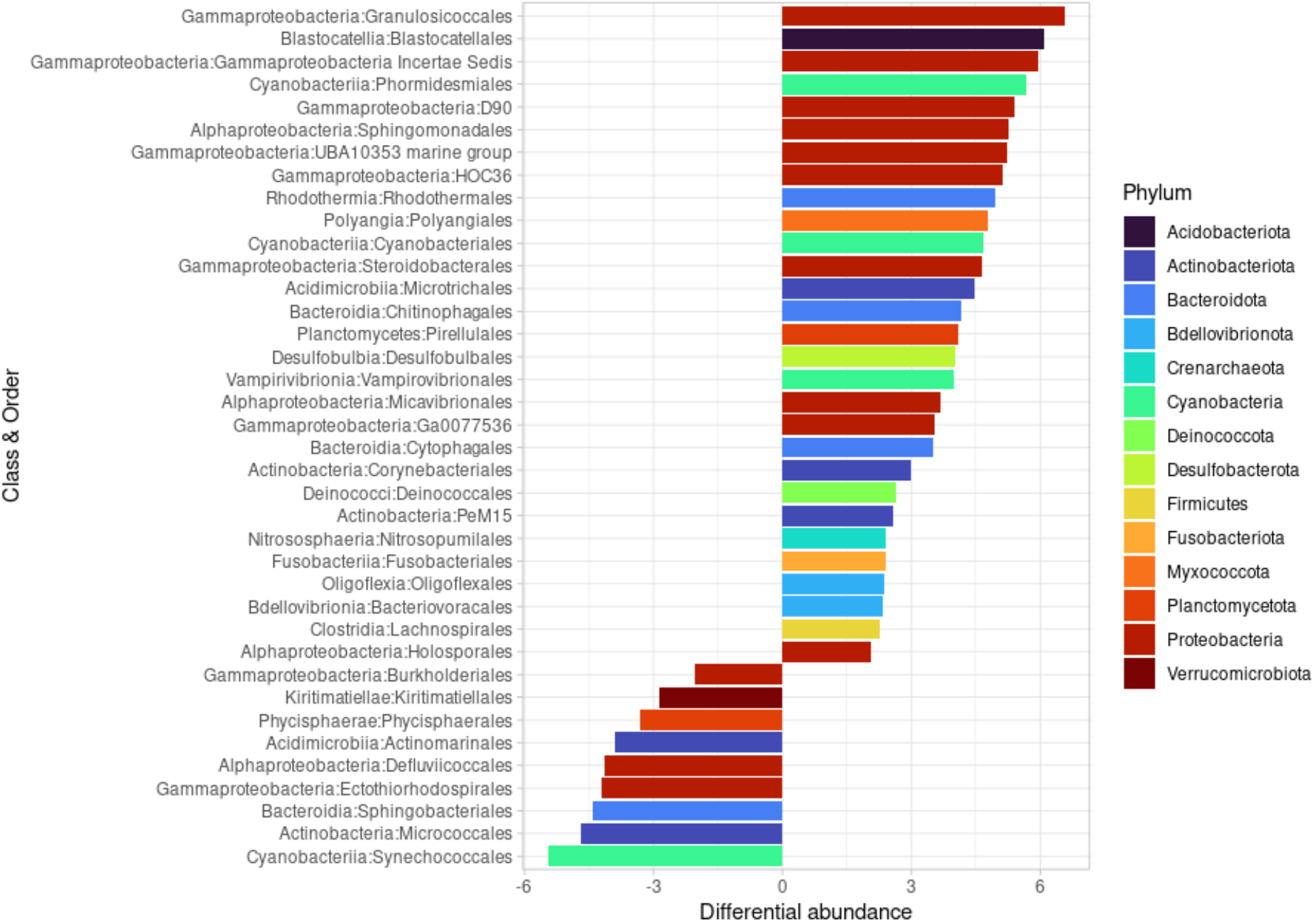
Differential taxonomic order abundance in leaf and water column samples. Positive values mean differential abundance in the leaves and negative values in the water column.

Among the predominant orders in the phylum, *Bacteroidota* (17%) was the order *Flavobacterales* (8%). They are also frequently found as early colonizers on marine surfaces and seagrasses^41,42^. In other studies, some photosynthetic and light-dependent members of *Bacteroidota* that harbor the nifH gene, e.g., *Chlorobaculum* and *Chlorobium* ^36^, were more abundant on leaves than in the water column. Other heterotrophic bacterial N_2_ fixers that may depend on seagrass photosynthetic exudates^36^ were found on *P. oceanica* leaves within the *Desulfobacterota* phylum. As part of the *P. oceanica* leaf microbiome, these groups are likely to collectively contribute to N_2_ fixation as a consortium of (directly or indirectly) light-dependent N_2_ fixers.

*Granulosicoccus* was the phylotype with the largest effect detected in the differential abundance analysis (Fig. 3). It has been often found as part of the phyllosphere microbiome of macroalgae and seagrasses^50–52^ having the potential for dissimilatory nitrate reduction to ammonium and the synthesis of vitamins that are needed by their macroalgae host ^51^. Among the potential denitrifiers, the gammaproteobacterium *Marinicella* was predominantly detected on *P. oceanica* leaves; it often contributes to denitrification in *Synechococcus*-dominated biofilms and anammox-concentrating reactors, ^53–55^.

*Planctomycetes* accounted for 2% of the microbial leaf community (Fig. 2) and were more abundant on the leaves than in the water column (Fig. 3). *Planctomycetes* are commonly found on macroalgae across the globe^56,57^ and can even dominate the *P. oceanica* leaf microbiome^35^. Members of this phylum have been linked to N_2_ fixation in surface ocean waters ^58^. Among *Planctomycetes* are also members that can utilize anammox to gain energy by anaerobically oxidizing NH_4_^+^ with NO_2_^-^ as the electron acceptoR^59,60^.

Finally, we found significantly higher relative abundances of the families Nitrosomonadaceae, Nitrospiraceae, Nitrospinaceae (AOB), and Nitrosopumilales (AOA) in the phyllosphere of *P. oceanica* (Suppl. Fig. 7), all of which include nitrifying members^18,61^. In particular, we found a higher relative abundance of *Nitrosopumilales* (family *Nitrosopumilaceae*) on leaves, which often show a higher affinity for ammonia than AOB^62,63^, further indicating that competition for NH_4_^+^ plays a major role on seagrass leaves.

### Ocean acidification accelerates N cycling towards higher N_2_ fixation and N uptake

Our results show that OA accelerated key N transformation processes associated with the phyllosphere of *P. oceanica*, while the prokaryotic community structure remained largely unaffected. To quantify N transformation rates under OA conditions, we incubated leaf sections from CO_2_ vents, where the plant and its epiphytic community are acclimated to long-term CO_2_ enrichment and lower pH. We found that daylight N_2_ fixation was significantly higher on leaves acclimated to low pH (Fig. 1a). The positive response of N_2_ fixation rates to elevated CO_2_ concentrations is supported by several studies with planktonic diazotrophs, such as *Trichodesmium, Crocosphaera*, and *Nodularia* (see review papers by ^20,64,65^). A widely accepted explanation for the positive influence of elevated CO_2_ concentrations on some diazotrophs is their ability to reallocate energy from the downregulation of carbon-concentrating mechanisms to N_2_ fixation^20,65^.

Notably, potential nitrification (PNR) was only detected under OA conditions in our incubations (Fig. 1b). Reduced pH is generally expected to negatively affect ammonium oxidation in the first step of nitrification^24,66^. However, some studies showed that increasing CO_2_ levels could lead to higher autotrophic nitrification rates by reducing CO_2_ limitation^21^ and that a diverse nitrifier community, such as that found in estuarine and coastal sediments, could adapt to a wider range of pH values^67^.

Ocean acidification is generally not expected to have a major, direct effect on denitrification and anammox, as both processes occur in anaerobic environments that already have elevated CO_2_ concentrations and low pH values^20,21^. However, on *P. oceanica* leaves under high CO_2_ conditions, an increase in both C^11^ and N_2_ fixation, as well as nitrification, may have favored the formation of anoxic microniches on the leaf biofilm and generated organic C and oxidized N compounds available for metabolism by denitrifying bacteria^21^.

We observed that NH_4_^+^ uptake rates were increased by 61 – 110% at the vent site and NO_3_^-^ uptake rates were increased by 52 - 201 % (Suppl. Fig. 5 c, d). At the ambient site, we measured higher epiphyte cover and lower net primary production and respiration^11^, which can affect nutrient uptake rates. Apostolaki et al.^68^ showed that N uptake in leaves decreases with increasing epiphyte load, suggesting that epiphyte overgrowth inhibits leaf N uptake in *P. oceanica*. On the other hand, the seagrass may adapt to an increased N demand due to higher productivity under OA. This agrees with Ravaglioli et al.^69^, who found overexpression of N transporter genes after nutrient addition at low pH, suggesting increased N uptake by the seagrass.

While N cycling on the *P. oceanica* phyllosphere accelerated under high CO_2_, the prokaryotic community structure remained largely unaffected. Similarly, Banister et al.^70^ found that the leaf-associated microbiome of the seagrass *Cymodocea nodosa* was stable across pH gradients at a comparable Mediterranean CO_2_ vent site. Conversely, colonization experiments using an inert substrate showed marked differences in coastal microbial biofilms between natural pH and vent-exposed sites^71^. A stable microbial community in our study supports the hypothesis of a microbiome that is regulated by interactions with its plant host^50^, while our biogeochemical measurements suggest the presence of coupled metabolisms between the seagrass and its microbiome contributing to plant health and adaptation in a high-CO_2_ world.

### Phyllosphere N cycling contributes to the holobiont N demand

We calculated daily rates in mmol N m^-2^ d^-1^ of plant and epiphyte-mediated N-cycling processes at vent and ambient pH based on a 12:12 light/dark cycle (Fig. 4 a, b).

**Fig. 4.**
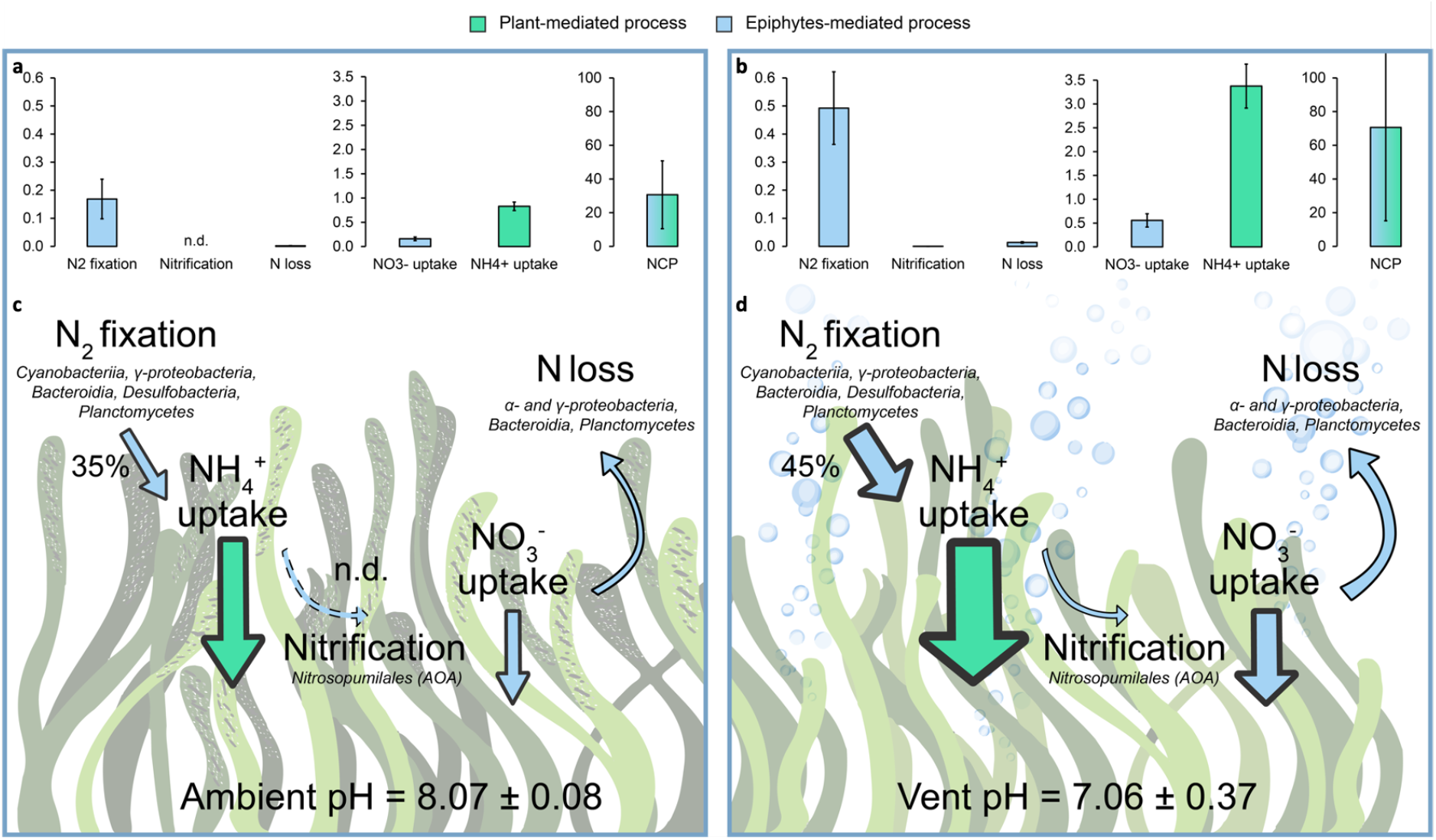
Overview of N cycling processes at (a, c) ambient and (b, d) vent pH. Panels a and b show daily metabolic rates (in mmol m^-2^ d^-1^) of plant- and epiphyte-mediated processes based on a 12:12 h light and dark cycle per area seagrass leaf. Panels c and d show the % contribution of N_2_ fixation to the estimated N demand of the plant based on NCP and C:N ratios and important taxa in the microbial community at ambient and vent pH. The arrow size indicates the magnitude of N cycling rates. Error bars indicate mean ± propagated standard errors of the mean.

We further used net community productivity (NCP) from Berlinghof et al.^11^ (using a photosynthetic quotient of 1), C:N ratios (Suppl. Fig. 6), average leaf density and dry weight per leaf at the ambient and vent site (Suppl. Table 4) to calculate the percentage of daily primary production of the seagrass holobiont (plant + epiphytes) that can be supported by leaf-associated N_2_ fixation (Fig. 4 c, d).

Although NCP, and thus the seagrass N demand, was higher under OA, the contribution of N_2_ fixation to meeting this demand was increased at vent pH. N_2_ fixation contributed with 169 ± 71 mmol N m^-2^ d^-1^ to 35 % of the seagrass N demand at ambient pH and with 493 ± 129 mmol N m^-2^ d^-1^ to 45 % at vent pH (Fig. 4). The contribution of N_2_ fixation to the seagrass N demand has been reported to be highly variable over seasonal^e.g.,36,72^ and spatial^36^ gradients. Integrating the seasonal values over a year, Agawin et al.^36^ calculated that ca. 15% of the annual plant N demand can be provided by aboveground N_2_ fixation in *P. oceanica* meadows. Further research (e.g., using NanoSIMS or longer-term incubations) should investigate how much of the N fixed by the epiphytic diazotrophs is actually transferred to the plant host.

A large fraction of the holobiont N demand was obtained through NH_4_^+^ uptake with 829 ± 87 mmol N m^-2^ d^-1^ at the ambient and 3376 ± 461 mmol N m^-2^ d^-1^ at the vent site (Fig. 4). NO_3_^-^ uptake, primarily attributed to the epiphytic community, contributed with 159 ± 37 mmol N m^-2^ d^-1^ at the ambient and 555 ± 139 mmol N m^-2^ d^-1^ at the vent site. NO_3_^-^ uptake rates were comparable to the annual average NO_3_^-^ leaf uptake by Lepoint et al.^37^ (1.2 g N m^-2^ yr^-1^ = 235 mmol N m^-2^ d^-1^). Conversely, NH_4_^+^ uptake rates were higher than their maximum values obtained in spring months (1300 μg N m^-2^ h^-1^ = 2227 mmol N m^-2^ d^-1^). However, Lepoint et al. also show that large seasonal differences can occur, with values ranging from 0 to 2227 mmol N m^-2^ d^-1 37^.

The total N gain (N_2_ fixation + NH_4_^+^ and NO_3_^-^ uptake - N loss) was 1115 ± 194 mmol N m^-2^ d^-1^ at the ambient and 4410 ± 727 mmol N m^-2^ d^-1^ at the vent site. Thus, OA tipped the balance decisively in favor of increased N gain. Taken together, our results show that major N cycling processes occur on *P. oceanica* leaves, and that epiphytes contribute to net N uptake by the holobiont. Ocean acidification accelerates N cycling, while the prokaryotic community structure remains largely unaffected. At a vent pH ∼ 7, high rates of microbial daylight N_2_ fixation on the phyllosphere of *P. oceanica* can partially sustain the increased C-fixation and thus N demand of the holobiont. Access to diverse N sources may help to avoid competition within the holobiont. Adaptation of marine plants to environmental changes is fundamental for their survival; here we show that functional plasticity of their N-cycling microbiome is a key factor in regulating seagrass holobiont functioning on a changing planet.

## Methods

### Study area and sampling

The study area is located at the islet of Castello Aragonese on the northeastern coast of the island of Ischia (Tyrrhenian Sea, Italy). This site is characterized by the presence of submarine CO_2_ vents of volcanic origin, which naturally generate a gradient in CO_2_ concentration and pH, without affecting the surrounding water temperature or salinity^73,74^. Around the islet, meadows of *P. oceanica* occur at depths of 0.5 - 3 m, also extending into vent zones with low pH. We selected two sites characterized by different pH regimes (Suppl. Table 1) at approximately 3 m water depth. The vent pH site was located in a vent area on the south side (40°43’50.5”N 13°57’47.2”E) and the ambient pH site was located on the north side of the bridge (40°43’54.8”N 13°57’47.1”E).

For the incubation experiments, shoots of *P. oceanica* were collected at each site on three days in September 2019 and transported directly to the laboratory. Sections of the central part of the leaf (3 cm in length) were cut off, selecting leaves with homogeneous epiphyte coverage, and avoiding heavily grazed and senescent parts of the plant, as described in Berlinghof et al.^11^ Epiphytes were carefully removed from half of the seagrass leaves with a scalpel, taking special care not to damage the plant tissue. Leaf sections from the vent pH and ambient pH sites, with epiphytes present (n = 4) or removed (n = 3), were used for dark and incubations.

Samples for microbial community analysis were collected in October 2019 at the vent and ambient site described above. Before disturbing the plants, we collected 5 L of seawater from the water column above the plants. Whole seagrass plants were collected, and the central part of the leaf was cut off with sterile tools, washed with sterile NaCl solution [0.8 % m/v] to remove loosely attached microorganisms, and transferred to 15 mL falcon tubes with sterile tweezers. The falcon tubes were kept in dry ice during transport to the laboratory (SZN Villa Dohrn, Ischia, Italy) and then stored at -20°C. In the laboratory, the seawater was immediately filtered on 0.2 μm cellulose nitrate membrane filters and the filters were stored at -20°C until further genetic analysis.

### Prokaryotic DNA extraction, amplification, and sequencing

DNA from seagrass and seawater samples was extracted using the Qiagen DNeasy Powersoil Kit (Qiagen). For seawater, the entire membrane filters were used, while for seagrass, we cut approximately 1 g of the central part of the leaf. Leaf samples were placed into 2 mL vials containing 600 μL of sterile NaCl solution [0.8 % m/v] and were vortexed three times for 30 s according to the protocol of the Seagrass Microbiome Project (https://seagrassmicrobiome.org). The solution was transferred to the Powerbead columns (Qiagen) and then processed according to the manufacturer’s instructions with slight modifications to increase DNA yield and quality, as described in Basili et al.^75^ The extracted DNA samples were quantified using a microvolume spectrophotometer (Thermo Scientific NanoDrop 2000c) and stored at -20 °C until processing.

Illumina MiSeq sequencing (2 × 300 bp paired-end protocol) of the hypervariable V4 region of the 16S rRNA gene was performed using the 515FB and 806RB bacteria- and archaea-specific primers^76^. The primers were removed from the raw sequence data using cutadapt v2.8^77^ and the fastq files were processed using the R package *DADA2*^78,79^. Quality filtering and denoising of the trimmed fastq files was performed using the following parameters: “truncLen = c(200, 200), maxEE = c(2, 2), truncQ = 2, ndmaxN = 0). Paired-end reads were then merged into amplicon sequence variants (ASVs); chimeric sequences were identified and removed. Prokaryotic taxonomy assignment was performed using the SILVA v138^80^ database. The complete pipeline is openly available in the research compendium accompanying this paper at https://github.com/luismmontilla/embrace. The sequences are available in the NCBI SRA database as the BioProject ID PRJNA824287.

### Bioinformatics and data analysis of the sequencing data

The ASV matrix was analyzed as a compositional dataset, as described in detail in other works^81,82^. Briefly, we transformed the raw pseudo-counts using the centered-log ratio to handle the data in a Euclidean space. We then tested the null hypothesis of no effect of the factors described above on the prokaryotic community associated with *P. oceanica* using a permutation-based multivariate analysis of variance derived from a Euclidean distance matrix. We performed this test using the *vegan* package for R^83,84^. In addition, we performed a differential abundance analysis of the ASVs using the ANOVA-like differential expression method implemented in the ALDEX2 package for R^85^. This algorithm produces consistent results, whereas other analyses can be variable depending on the parameters set by the researcher or required by the dataset^86^.

### Dinitrogen fixation rate measurements

The ^15^N_2_-enriched seawater addition method was used to determine N_2_ fixation rates^87^. Stock solutions of 0.22 μm filtered and ^15^N_2_-enriched water from the two study sites (vent and ambient pH) were prepared and gently transferred to 24 mL glass vials to minimize gas exchange with the atmosphere. Subsequently, seagrass leaves with (n=4) and without epiphytes (n=3) were added and the vials were sealed without leaving any headspace. Additionally, vials with 0.22 μm filtered but unenriched site water containing leaves with epiphytes served as controls to account for any natural increase in δ^15^N (n=3). The vials were incubated on a shaker (Stuart Orbital Shaker SSL1; 30 rpm); vials for dark incubations were covered with aluminum foil. Incubations were performed in a temperature-controlled room at 22°C. After an incubation period of T0 = 0 h, T1 = 5 h, and T2 = 9 h light/ 8 h dark, three or four vials from each treatment were opened for sampling. At the beginning and end of the incubation, oxygen concentrations in the incubation vials were measured using a fiber-optic oxygen sensor (FireStingO2, PyroScience), and pH was measured using a pH meter (Multi 3430, WTW).

For tissue analysis, epiphytes and seagrass leaves were transferred separately into Eppendorf tubes and freeze-dried for 72 h. They were then homogenized in a mortar and weighed in tin cups to determine carbon (%C) and nitrogen content (%N), and ^15^N incorporation. Water samples were transferred to 12 mL exetainers (Labco Ltd) and fixed with 200 μL of 7 M ZnCl_2_ for ^29^N_2_ and ^30^N_2_ analyses to calculate atom% excess of the medium. In addition, samples for the analysis of dissolved inorganic nitrogen (DIN: NH_4_^+^, NO^2-^, NO_x_^-^) and PO_4_^3-^ were transferred to 20 ml HDPE vials and stored at -20°C until further analysis.

Carbon (%C) and nitrogen (%N) content and the isotopic composition (δ^13^C, δ^15^N) in seagrass leaves and epiphyte tissue were analyzed by isotope ratio mass spectrometry (IRMS, Delta plus V, Thermo Scientific) coupled to an elemental analyzer (Flash EA1112, Thermo Scientific) at Aarhus University (Denmark). ^15^N_2_ fixation rates were calculated according to Montoya et al.^88^:

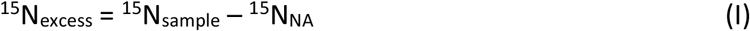

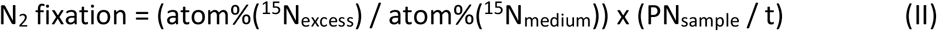

^15^N_sample_ is the ^15^N content of the samples after exposure to ^15^N_2_ enriched seawater, and ^15^N_NA_ is the ^15^N content in natural abundance samples without ^15^N_2_ exposure. The enrichment of samples (^15^N_excess_) was considered significant for samples with a value greater than 2.5 times the standard deviation of the mean of the natural abundance samples. ^15^N_medium_ is the enrichment of the incubation medium at the end of the incubations. With our approach, we achieved an enrichment of ∼16.0 atom %^15^N in the incubation vials. PN_sample_ is the N content of the sample (μg), and t represents the incubation time (h). ^15^N_2_ fixation rates were normalized per seagrass leaf area (cm^2^). The C:N molar ratio was determined as: C:N= (% C/12) / (% N/14).

Dissolved nutrient concentrations (NH_4_^+^, NO_2_^-^, NO_x_^-^, PO_4_^-^) were measured with a continuous flow analyzer (Flowsys, SYSTEA S.p.A.). NO_3_^-^ concentrations were calculated as the difference between NO_x_^-^ and NO_2_^-^. Subsequently, nutrient fluxes were calculated as the difference between final and initial nutrient concentrations, corrected for controls, and normalized to leaf area.

### Potential Nitrification rate measurements

Nitrification potential was determined using stock solutions of 0.22 μm filtered water from the study sites (vent and ambient pH site) with an ambient NH_4_^+^ concentration of 0.65 μM that was enriched with ^15^NH_4_^+^ (≥98 atom %^15^N) to a final concentration of 20 μM. The incubation was performed as described above with sampling times at T0 = 0h, T1 = 2h, T2 = 5h, and T3 = 9 h light/ 8 h dark. Water samples were filtered at 0.22 μm, transferred to 15 mL polypropylene tubes, and stored at -20°C for the analysis of NO_3_^-^ production. Vials with 0.22 μm filtered site water without the addition of ^15^NH_4_^+^ and without leaves served as controls for background microbial activity in the water column (n=3).

Isotopic samples for ^15^NO_3_^-^ production were analyzed by isotope ratio mass spectrometry (IRMS) using a modified version of the Ti(III) reduction method described by Altabet et al.^89^ Sample aliquots for nitrification analysis (3 mL) were acidified by adding 10 μL of 2.5 nM sulfanilic acid in 10% HCl to each 1 mL of sample, then added to 3 mL of the international standard USGS-32 (δ^15^N = +180‰) in a 12 mL exetainer, so that the final concentration of USGS-32 was 0.1 ppm NO_3_-N (∼7 μM NO_3_^-^). After combining the sample with the standard, the exetainer headspace was flushed with argon for 2 minutes. NO_3_^-^ was then converted to nitrous oxide (N_2_O) for stable N isotope analysis by adding 200 μL zinc-treated 30% TiCl_3_. The exetainers were immediately sealed with a gas-tight, pierceable, chlorobutyl rubber septum and the final reaction volume was 6.15 mL. The Ti(III)-treated samples were left at room temperature for >12 h to convert NO_3_^-^ to N_2_O. The headspace of the exetainer was sampled with a double-holed needle using a CTC PAL autosampler and a modified flush-fill line of a GasBench device (Thermo Scientific). The flush rate was ca. 25 mL min^-1^ and the flushing time was 5.5 min. The headspace sample was passed through a magnesium perchlorate and ascarite trap to remove water and CO_2_, respectively, and then collected in a sample loop (50 cm PoraPlot Q; ø = 0.53 mm; Restek) submersed in liquid nitrogen. N_2_O in the sample was then separated from CO_2_ and other gases by injecting onto a Carboxen 1010 PLOT column (30 m × 0.53 mm, 30 μm film thickness, Supelco; temp = 90 °C, flow rate 2.6 mL min^-1^) with helium as carrier gas. The sample was then transferred to a MAT253 PLUS IRMS via a Conflo interface (ThermoScientific). δ^15^N values were determined relative to the N_2_O working gas, and then corrected for linearity according to the peak height relationship and the titanium-to-sample ratio^89^; the absolute value of the linear correction term was <1.3‰ for all samples. The corrected values were then normalized to the δ^15^N-air scale by simultaneous analysis of the international standards USGS32, USGS34, and USGS35. The δ^15^N value of NO_3_^-^ in the sample was finally determined via a mass balance of the relative NO_3_^-^ concentrations of the sample and USGS32, the measured δ^15^N value of the mixture, and the accepted δ^15^N value of USGS32. The external precision of the δ^15^N measurement (± one standard deviation of the mean) determined for an in-house standard was 1.1‰.

Potential nitrification rates (PNR) were calculated using an equation modified from Beman et al.^24^:

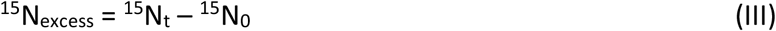

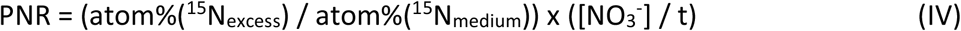

^15^N_t_ is the ^15^N content of the samples in the NO_3_^−^ pool measured at time t, and ^15^N_0_ is the ^15^N content in the NO_3_^−^ pool measured at the beginning of the incubations. The enrichment of samples (^15^N_excess_) was considered significant for samples with a value greater than 2.5 times the standard deviation of the mean of the T_0_ samples. ^15^N_medium_ is the enrichment of the incubation medium at the end of the incubations. Based on the NH_4_^+^ concentrations measured before and after the addition of ^15^NH_4_^+^, this resulted in a theoretical enrichment of ∼95.9 atom %^15^N in the incubation medium. [NO_3_^-^] is the concentration of NO_3_^-^ (μM) and t is the incubation time (h). Potential nitrification rates were normalized per seagrass leaf area (cm^2^) and corrected for the rates in control incubations without organisms.

### Potential Anammox and Denitrification rate measurements

To determine the rates of N loss via N_2_ production (combined denitrification and anammox), stock solutions of 0.22 μm filtered water from the two study sites (vent and ambient pH) with an ambient NO_3_^-^ concentration of 1.94 μM were enriched with ^15^NO_3_^-^ (≥98 atom %^15^N) to a final concentration of 10 μM. The incubation was performed as in the N_2_ fixation experiment (see “*Dinitrogen fixation rate measurements”*), with sampling times at T0 = 0 h, T1 = 2 h, T2 = 5 h, and T3 = 9 h light/ 8h dark. Vials with 0.22 μm filtered site water without the addition of NO_3_^-^ and without leaves served as controls for background microbial activity in the water column (n=3). Water samples were transferred into 12 mL exetainers and fixed with 200 μL of 7 M ZnCl_2_ for ^29^N_2_ and ^30^N_2_ analyses.

Isotopic samples for ^29^N_2_ and ^30^N_2_ production were analyzed by gas chromatography-isotope ratio mass spectrometry (GasBench, Thermo Scientific). ^29^N_2_ and ^30^N_2_ concentrations were calculated via linear regression of a standard curve with N_2_ air standards. Production rates of ^15^N-enriched N_2_ gas were calculated from the difference in ^29^N_2_ or ^30^N_2_ concentrations between T1 (2 h) and T2 (5 h), as we observed a lag phase from T0 to T1. Because the changes in ^29^N_2_ and ^30^N_2_ concentrations were very small (Suppl. Table 5), we decided to report ^29^N_2_ and ^30^N_2_ production rates instead of further transforming the data to calculate denitrification or anammox rates. ^29^N_2_ and ^30^N_2_ production rates were normalized to seagrass leaf area (cm^2^) and corrected for the rates in control incubations without organisms.

### Data analysis of the biogeochemistry data

We tested for normality and homogeneity of variances before each analysis using Shapiro-Wilk’s and Levene’s tests and transformed data or removed outliers if normality and homogeneity of variances were not met. We tested the effects of pH (vent pH vs. ambient pH), treatment (with and without epiphytes), and their interaction on the ^15^N_2_ incorporation rates, potential nitrification rates (PNR), ^29^N_2_ and ^30^N_2_ production rates, and the nutrient fluxes using two-way ANOVAs (type II). We tested the effects of pH (vent pH vs. ambient pH) on the C:N ratios of leaves and epiphytes using a one-way ANOVA (type II). All statistical analyses were performed with R^78^ (version 4.1.2) using the packages *car* and *emmeans*.

## Data availability

All data needed to evaluate the conclusions of the paper are provided in the paper and/or in the Supplementary Materials. Source data for all N isotope experiments, net dissolved inorganic N fluxes and C:N ratios, and 16S rRNA gene ASV tables, are provided with this paper. Raw sequencing data supporting the results of this study have been deposited in the NCBI GenBank database (https://www.ncbi.nlm.nih.gov/bioproject/) with the accession code: BioProject PRJNA824287. The source data are provided with this paper.

## Acknowledgements

We thank L. Polaková and K. Umbria-Salinas of the SoWa Stable Isotope Facility (České Budějovice, CZ) for their assistance with nitrate isotope measurements. This research was supported by a Ph.D. fellowship co-funded by the Stazione Zoologica Anton Dohrn (SZN) and the University of Bremen (to J.B. and F.P.), a Ph.D. fellowship funded by the Open University – SZN Ph.D. Program (to L.M.M.), and a SZN postdoctoral fellowship (to U.M.).

## Author contributions

F.P., G.M.Q., U.M., C.W. and U.C. designed the study. F.P., G.M.Q., U.M. and U.C. performed the experiments. J.B., U.M. and T.B.M. performed the mass spectrometry analyses. L.M.M. and G.M.Q. performed the molecular analyses. F.M. and M.A. performed the nutrient analyses. J.B. and L.M.M. analyzed the results. J.B., L.M.M. and U.C. wrote the manuscript with contributions from all co-authors.

## References

1. Hemminga, M. A. & Duarte, C. M. Seagrass Ecology. Seagrass Ecology (Cambridge University Press, 2000). doi:10.1017/cbo9780511525551.

2. Björk, M., Short, F. T., Mcleod, E. & Beer, S. Managing seagrasses for resilience to climate change. (IUCN, Gland, Switzerland, 2008).

3. Hyman, A. C., Frazer, T. K., Jacoby, C. A., Frost, J. R. & Kowalewski, M. Long-term persistence of structured habitats: seagrass meadows as enduring hotspots of biodiversity and faunal stability. Proc. R. Soc. B 286, (2019).

4. Fourqurean, J. W. et al. Seagrass ecosystems as a globally significant carbon stock. Nat Geosci 5, 505–509 (2012).

5. Duarte, C. M., Kennedy, H., Marbà, N. & Hendriks, I. Assessing the capacity of seagrass meadows for carbon burial: Current limitations and future strategies. Ocean Coast Manag 83, 32–38 (2013).

6. Hendriks, I. E. et al. Photosynthetic activity buffers ocean acidification in seagrass meadows. Biogeosciences 11, 333–346 (2014).

7. Lacoue-Labarthe, T. et al. Impacts of ocean acidification in a warming Mediterranean Sea: An overview. Reg Stud Mar Sci 5, 1–11 (2016).

8. Koch, M., Bowes, G., Ross, C. & Zhang, X.-H. Climate change and ocean acidification effects on seagrasses and marine macroalgae. Glob Chang Biol 19, 103–132 (2013).

9. Cox, T. E. et al. Effects of ocean acidification on Posidonia oceanica epiphytic community and shoot productivity. Journal of Ecology 103, 1594–1609 (2015).

10. Hernán, G. et al. Seagrass (Posidonia oceanica) seedlings in a high-CO 2 world: from physiology to herbivory. Sci Rep 6, (2016).

11. Berlinghof, J. et al. The role of epiphytes in seagrass productivity under ocean acidification. Sci Rep 12, (2022).

12. Cox, T. E. et al. Effects of in situ CO2 enrichment on structural characteristics, photosynthesis, and growth of the Mediterranean seagrass Posidonia oceanica. Biogeosciences 13, 2179–2194 (2016).

13. Scartazza, A. et al. Carbon and nitrogen allocation strategy in Posidonia oceanica is altered by seawater acidification. Science of the Total Environment 607, 954–964 (2017).

14. Donnarumma, L., Lombardi, C., Cocito, S. & Gambi, M. C. Settlement pattern of Posidonia oceanica epibionts along a gradient of ocean acidification: an approach with mimics. Mediterr Mar Sci 15, 498–509 (2014).

15. Mecca, S., Casoli, E., Ardizzone, G. & Gambi, M. C. Effects of ocean acidification on phenology and epiphytes of the seagrass Posidonia oceanica at two CO2 vent systems of Ischia (Italy). Mediterr Mar Sci 21, 70–83 (2020).

16. Gravili, C., Cozzoli, F. & Gambi, M. C. Epiphytic hydroids on Posidonia oceanica seagrass meadows are winner organisms under future ocean acidification conditions: evidence from a CO2 vent system (Ischia Island, Italy). European Zoological Journal 88, 472–486 (2021).

17. Hemminga, M. A., Harrison, P. G. & Van Lent, F. The balance of nutrient losses and gains in seagrass meadows. Mar Ecol Prog Ser 71, 85–96 (1991).

18. Kuypers, M. M. M., Marchant, H. K. & Kartal, B. The microbial nitrogen-cycling network. Microbial Biogeochemistry 16, 263–276 (2018).

19. Wyatt, N. J. et al. Effects of high CO2 on the fixed nitrogen inventory of the Western English Channel. J Plankton Res 32, 631–641 (2010).

20. Wannicke, N., Frey, C., Law, C. S. & Voss, M. The response of the marine nitrogen cycle to ocean acidification. Glob Chang Biol 24, 5031–5043 (2018).

21. Hutchins, D. A., Mulholland, M. R. & Fu, F. Nutrient Cycles and Marine Microbes in a CO_2_-Enriched Ocean. Oceanography 22, 128–145 (2009).

22. Levitan, O. et al. Elevated CO2 enhances nitrogen fixation and growth in the marine cyanobacterium Trichodesmium. Glob Chang Biol 13, 531–538 (2007).

23. Kranz, S. A. et al. Combined Effects of CO 2 and Light on the N2-Fixing Cyanobacterium Trichodesmium IMS101: Physiological Responses. Plant Physiol 154, 334–345 (2010).

24. Beman, J. M. et al. Global declines in oceanic nitrification rates as a consequence of ocean acidification. PNAS 108, 208–213 (2011).

25. Ugarelli, K., Chakrabarti, S., Laas, P. & Stingl, U. The Seagrass Holobiont and Its Microbiome. Microorganisms 5, (2017).

26. Tarquinio, F., Hyndes, G. A., Laverock, B., Koenders, A. & Säwström, C. The seagrass holobiont: Understanding seagrass-bacteria interactions and their role in seagrass ecosystem functioning. FEMS Microbiol Lett 366, 1–15 (2019).

27. Mohr, W. et al. Terrestrial-type nitrogen-fixing symbiosis between seagrass and a marine bacterium. Nature 2021 1–5 (2021) doi:10.1038/s41586-021-04063-4.

28. Agawin, N. S. R. et al. Significant nitrogen fixation activity associated with the phyllosphere of Mediterranean seagrass Posidonia oceanica: first report. Mar Ecol Prog Ser 551, 53–62 (2016).

29. Garcias-Bonet, N., Arrieta, J. M., Duarte, C. M. & Marbà, N. Nitrogen-fixing bacteria in Mediterranean seagrass (Posidonia oceanica) roots. Aquat Bot 131, 57–60 (2016).

30. Lehnen, N. et al. High rates of microbial dinitrogen fixation and sulfate reduction associated with the Mediterranean seagrass Posidonia oceanica. Syst Appl Microbiol 39, 476–483 (2016).

31. Agawin, N. S. R., Ferriol, P. & Sintes, E. Simultaneous measurements of nitrogen fixation in different plant tissues of the seagrass Posidonia oceanica. Mar Ecol Prog Ser 611, 111–127 (2019).

32. Ling, J. et al. Community Composition and Transcriptional Activity of Ammonia-Oxidizing Prokaryotes of Seagrass Thalassia hemprichii in Coral Reef Ecosystems. Front. Microbiol 9, 7 (2018).

33. Noisette, F., Depetris, A., Kühl, M. & Brodersen, K. E. Flow and epiphyte growth effects on the thermal, optical and chemical microenvironment in the leaf phyllosphere of seagrass (Zostera marina). J R Soc Interface 17, (2020).

34. Brodersen, K. E. & Kühl, M. Effects of Epiphytes on the Seagrass Phyllosphere. Front Mar Sci 9, 1–10 (2022).

35. Kohn, T. et al. The Microbiome of Posidonia oceanica Seagrass Leaves Can Be Dominated by Planctomycetes. Front. Microbiol 11, 1458 (2020).

36. Agawin, N. S. R., Ferriol, P., Sintes, E. & Moyà, G. Temporal and spatial variability of in situ nitrogen fixation activities associated with the Mediterranean seagrass Posidonia oceanica meadows. Limnol Oceanogr 62, 2575–2592 (2017).

37. Lepoint, G., Millet, S., Dauby, P., Gobert, S. & Bouquegneau, J. M. Annual nitrogen budget of the seagrass Posidonia oceanica as determined by in situ uptake experiments. Mar Ecol Prog Ser 237, 87–96 (2002).

38. Salk, K. R., Erler, D. V., Eyre, B. D., Carlson-Perret, N. & Ostrom, N. E. Unexpectedly high degree of anammox and DNRA in seagrass sediments: Description and application of a revised isotope pairing technique. Geochim Cosmochim Acta 211, 64–78 (2017).

39. Touchette, B. W. & Burkholder, J. A. M. Review of nitrogen and phosphorus metabolism in seagrasses. J Exp Mar Biol Ecol 250, 133–167 (2000).

40. Dang, H., Li, T., Chen, M. & Huang, G. Cross-ocean distribution of Rhodobacterales bacteria as primary surface colonizers in temperate coastal marine waters. Appl Environ Microbiol 74, 52–60 (2008).

41. Mejia, A. Y. et al. Assessing the ecological status of seagrasses using morphology, biochemical descriptors and microbial community analyses. A study in Halophila stipulacea (Forsk.) Aschers meadows in the northern Red Sea. Ecol Indic 60, 1150–1163 (2016).

42. Trevathan-Tackett, S. M. et al. Spatial variation of bacterial and fungal communities of estuarine seagrass leaf microbiomes. Aquatic Microbial Ecology 84, 59–74 (2020).

43. Li, Y. et al. Microbiota and functional analyses of nitrogen-fixing bacteria in root-knot nematode parasitism of plants. Microbiome 11, 1–23 (2023).

44. Lesser, M. P., Morrow, K. M., Pankey, S. M. & Noonan, S. H. C. Diazotroph diversity and nitrogen fixation in the coral Stylophora pistillata from the Great Barrier Reef. ISME J 12, 813–824 (2018).

45. Moynihan, M. A. et al. Coral-associated nitrogen fixation rates and diazotrophic diversity on a nutrient-replete equatorial reef. ISME J 16, 233–246 (2022).

46. Lindström, K. & Mousavi, S. A. Minireview Effectiveness of nitrogen fixation in rhizobia. Microb Biotechnol 13, 1314–1335 (2020).

47. Avis, T. J., Gravel, V., Antoun, H. & Tweddell, R. J. Multifaceted beneficial effects of rhizosphere microorganisms on plant health and productivity. Soil Biol Biochem 40, 1733–1740 (2008).

48. Bergman, B., Sandh, G., Lin, S., Larsson, J. & Carpenter, E. J. Trichodesmium – a widespread marine cyanobacterium with unusual nitrogen fixation properties. FEMS Microbiol Rev 37, 286–302 (2013).

49. Berrendero, E. et al. Nitrogen fixation in a non-heterocystous cyanobacterial mat from a mountain river. Sci Rep 6, 30920 (2016).

50. Crump, B. C., Wojahn, J. M., Tomas, F. & Mueller, R. S. Metatranscriptomics and amplicon sequencing reveal mutualisms in seagrass microbiomes. Front Microbiol (2018) doi:10.3389/fmicb.2018.00388.

51. Weigel, B. L., Miranda, K. K., Fogarty, E. C., Watson, A. R. & Pfister, C. A. Functional Insights into the Kelp Microbiome from Metagenome-Assembled Genomes. mSystems 7, (2022).

52. Sanders-Smith, R. et al. Host-Specificity and Core Taxa of Seagrass Leaf Microbiome Identified Across Tissue Age and Geographical Regions. Front Ecol Evol 8, 1–13 (2020).

53. Zhang, Z. et al. Long-Term Survival of Synechococcus and Heterotrophic Bacteria without External Nutrient Supply after Changes in Their Relationship from Antagonism to Mutualism. mBio 12, (2021).

54. Van Duc, L. et al. High growth potential and nitrogen removal performance of marine anammox bacteria in shrimp-aquaculture sediment. Chemosphere 196, 69–77 (2018).

55. Yin, S., Li, J., Dong, H. & Qiang, Z. Unraveling the nitrogen removal properties and microbial characterization of “Candidatus Scalindua”-dominated consortia treating seawater-based wastewater. Science of The Total Environment 786, 147470 (2021).

56. Bondoso, J. et al. Epiphytic Planctomycetes communities associated with three main groups of macroalgae. FEMS Microbiol Ecol 93, (2017).

57. Lage, O. M., Bondoso, J., Luis, R., Comolli, L. & Bengtsson, M. Planctomycetes and macroalgae, a striking association. (2014) doi:10.3389/fmicb.2014.00267.

58. Delmont, T. O. et al. Nitrogen-fixing populations of Planctomycetes and Proteobacteria are abundant in surface ocean metagenomes. Nat Microbiol 3, 804–813 (2018).

59. Strous, M. et al. Missing lithotroph identified as new planctomycete. Letters to Nature 400, 446–449 (1999).

60. Jetten, M. S. M. et al. Biochemistry and molecular biology of anammox bacteria. Crit Rev Biochem Mol Biol 44, 65–84 (2009).

61. Hutchins, D. A. & Capone, D. G. The marine nitrogen cycle: new developments and global change. Nat Rev Microbiol 20, 401–414 (2022).

62. Jung, M.-Y. et al. Ammonia-oxidizing archaea possess a wide range of cellular ammonia affinities. ISME 16, 272–283 (2022).

63. Martens-Habbena, W., Berube, P. M., Urakawa, H., De La Torre, J. R. & Stahl, D. A. Ammonia oxidation kinetics determine niche separation of nitrifying Archaea and Bacteria. Nature Letters 461, 976–981 (2009).

64. Liu, J., Weinbauer, M. G., Maier, C., Dai, M. & Gattuso, J. P. Effect of ocean acidification on microbial diversity and on microbe-driven biogeochemistry and ecosystem functioning. Aquatic Microbial Ecology 61, 291–305 (2010).

65. Kroeker, K. J. et al. Impacts of ocean acidification on marine organisms: quantifying sensitivities and interaction with warming. Glob Chang Biol 19, 1884–1896 (2013).

66. Kitidis, V. et al. Impact of ocean acidification on benthic and water column ammonia oxidation. Geophys Res Lett 38, (2011).

67. Fulweiler, R. W., Emery, H. E., Heiss, E. M. & Berounsky, V. M. Assessing the Role of pH in Determining Water Column Nitrification Rates in a Coastal System. Estuaries and Coasts 34, 1095–1102 (2011).

68. Apostolaki, E. T., Vizzini, S. & Karakassis, I. Leaf vs. epiphyte nitrogen uptake in a nutrient enriched Mediterranean seagrass (Posidonia oceanica) meadow. Aquat Bot 96, 58–62 (2012).

69. Ravaglioli, C. et al. Nutrient Loading Fosters Seagrass Productivity Under Ocean Acidification. Sci Rep 7, (2017).

70. Banister, R. B., Schwarz, M. T., Fine, M., Ritchie, K. B. & Muller, E. M. Instability and Stasis Among the Microbiome of Seagrass Leaves, Roots and Rhizomes, and Nearby Sediments Within a Natural pH Gradient. Microb Ecol (2021) doi:10.1007/s00248-021-01867-9.

71. Lidbury, I., Johnson, V., Hall-Spencer, J. M., Munn, C. B. & Cunliffe, M. Community-level response of coastal microbial biofilms to ocean acidification in a natural carbon dioxide vent ecosystem. Mar Pollut Bull (2012) doi:10.1016/j.marpolbul.2012.02.011.

72. Cardini, U., Van Hoytema, N., Bednarz, V. N., Al-Rshaidat, M. M. D. & Wild, C. N2 fixation and primary productivity in a red sea Halophila stipulacea meadow exposed to seasonality. Limnol. Oceanogr 63, 786–798 (2018).

73. Hall-Spencer, J. M. et al. Volcanic carbon dioxide vents show ecosystem effects of ocean acidification. Nature 454, (2008).

74. Foo, S. A., Byrne, M., Ricevuto, E. & Gambi, M. C. The carbon dioxide vents of Ischia, Italy, a natural system to assess impacts of ocean acidification on marine ecosystems: An overview of research and comparisons with other vent systems. Oceanography and Marine Biology 56, 237–310 (2018).

75. Basili, M. et al. Major Role of Surrounding Environment in Shaping Biofilm Community Composition on Marine Plastic Debris. Front Mar Sci 7, (2020).

76. Walters, W. et al. Improved Bacterial 16S rRNA Gene (V4 and V4-5) and Fungal Internal Transcribed Spacer Marker Gene Primers for Microbial Community Surveys. mSystems 1, (2016).

77. Martin, M. Cutadapt removes adapter sequences from high-throughput sequencing reads. EMBnet J 17, 10 (2011).

78. R Core team. R: A language environment for statistical computing. Preprint at (2021).

79. Callahan, B. J. et al. DADA2: High-resolution sample inference from Illumina amplicon data. Nat Methods 13, 581–583 (2016).

80. Quast, C. et al. The SILVA ribosomal RNA gene database project: improved data processing and web-based tools. Nucleic Acids Res 41, D590–D596 (2012).

81. Quinn, T. P., Erb, I., Richardson, M. F. & Crowley, T. M. Understanding sequencing data as compositions: An outlook and review. Bioinformatics (2018) doi:10.1093/bioinformatics/bty175.

82. Gloor, G. B., Macklaim, J. M., Pawlowsky-Glahn, V. & Egozcue, J. J. Microbiome Datasets Are Compositional: And This Is Not Optional. Front Microbiol 8, (2017).

83. Oksanen, J. et al. vegan: Community ecology package. Preprint at (2020).

84. Anderson, M. J. A new method for non-parametric multivariate analysis of variance. Austral Ecol 26, 32–46 (2001).

85. Fernandes, A. D., Macklaim, J. M., Linn, T. G., Reid, G. & Gloor, G. B. ANOVA-Like Differential Expression (ALDEx) Analysis for Mixed Population RNA-Seq. PLoS One 8, e67019 (2013).

86. Nearing, J. T. et al. Microbiome differential abundance methods produce different results across 38 datasets. Nat Commun 13, 342 (2022).

87. Klawonn, I. et al. Simple approach for the preparation of 15−15 N 2-enriched water for nitrogen fixation assessments: evaluation, application and recommendations. Front Microbiol 6, (2015).

88. Montoya, J. P., Voss, M., Kähler, P. & Capone, D. G. A Simple, High-Precision, High-Sensitivity Tracer Assay for N2 Fixation. Appl Environ Microbiol 62, 986–993 (1996).

89. Altabet, M. A., Wassenaar, L. I., Douence, C. & Roy, R. A Ti(III) reduction method for one-step conversion of seawater and freshwater nitrate into N2O for stable isotopic analysis of 15N/14N, 18O/16O and 17O/16O. Rapid Communications in Mass Spectrometry 33, 1227–1239 (2019).

