## Supplementary material for "Accelerated Nitrogen Cycling on Seagrass Leaves in a High-CO_2_ World"

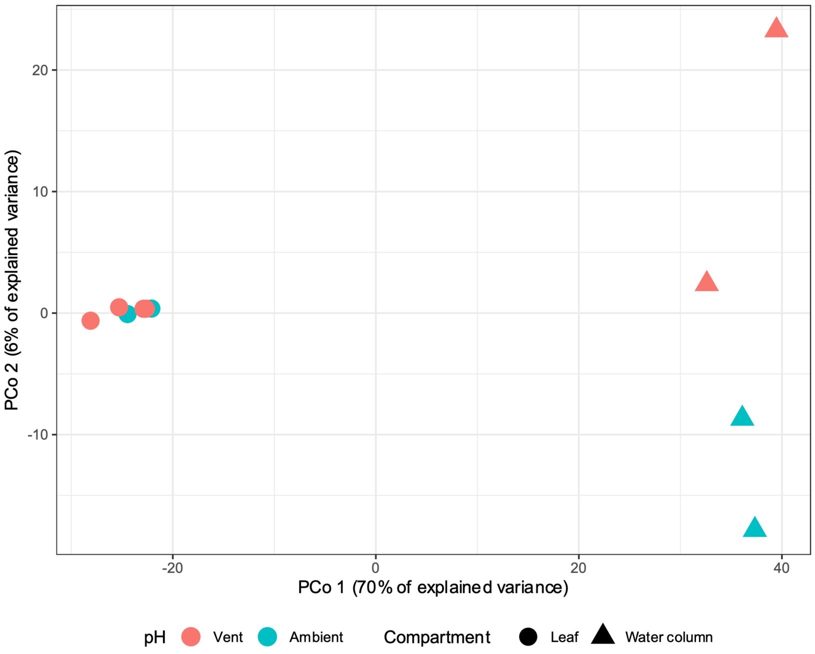


**Suppl. Figure 1.** Principal coordinates analysis of the prokaryote community from the leaves and water column on both pH regimes.

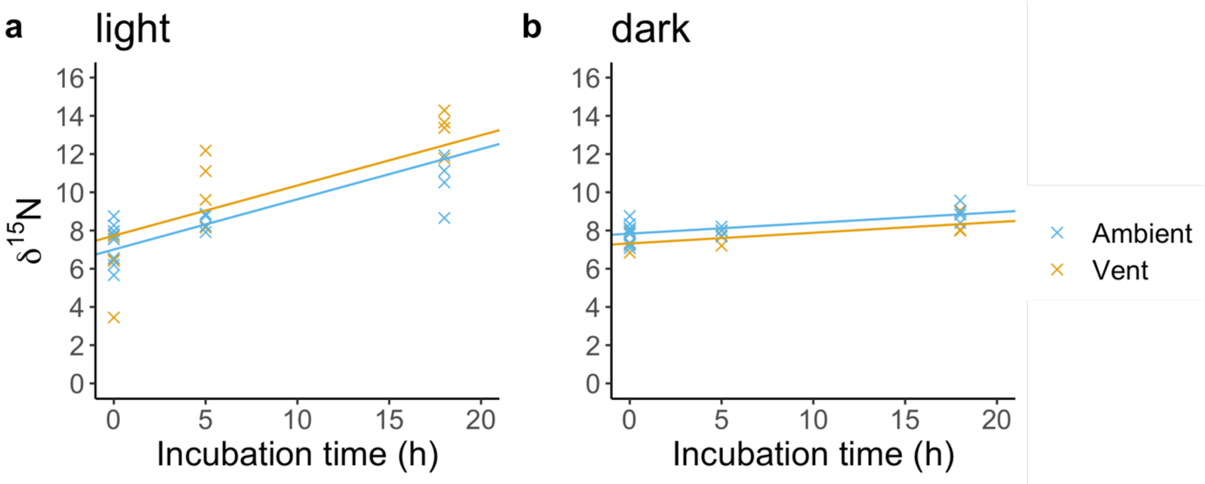


**Suppl. Figure 2.** ™15N increase during light (**a**) and dark (**b**) incubations in epiphytes from the ambient and the vent site. Solid lines represent linear regressions.

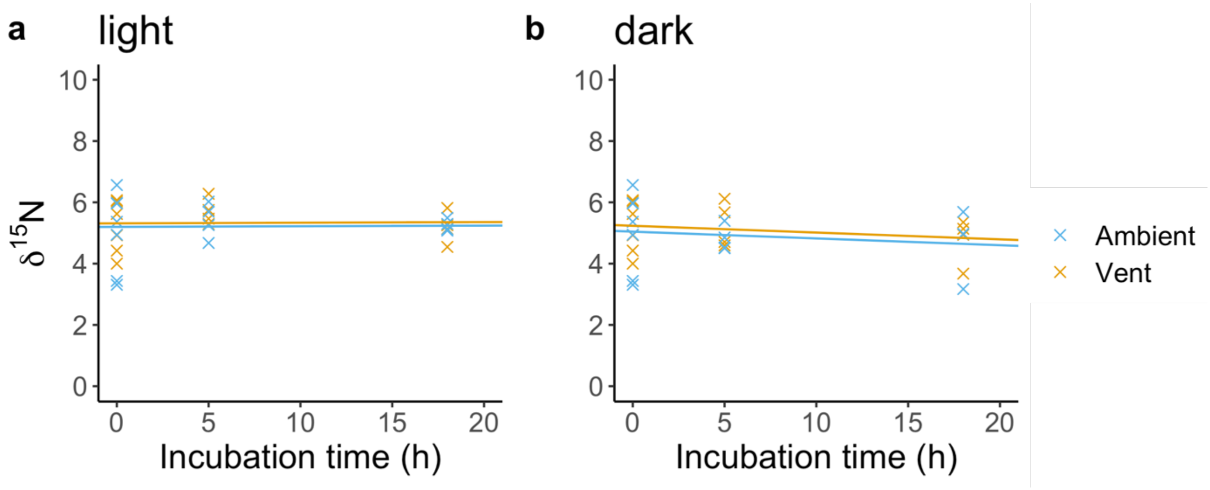


**Suppl. Figure 3**. ™$^{15}$N increase during light (**a**) and dark (**b**) incubations in seagrass leaf sections from the ambient and the vent site. Solid lines represent linear regressions.

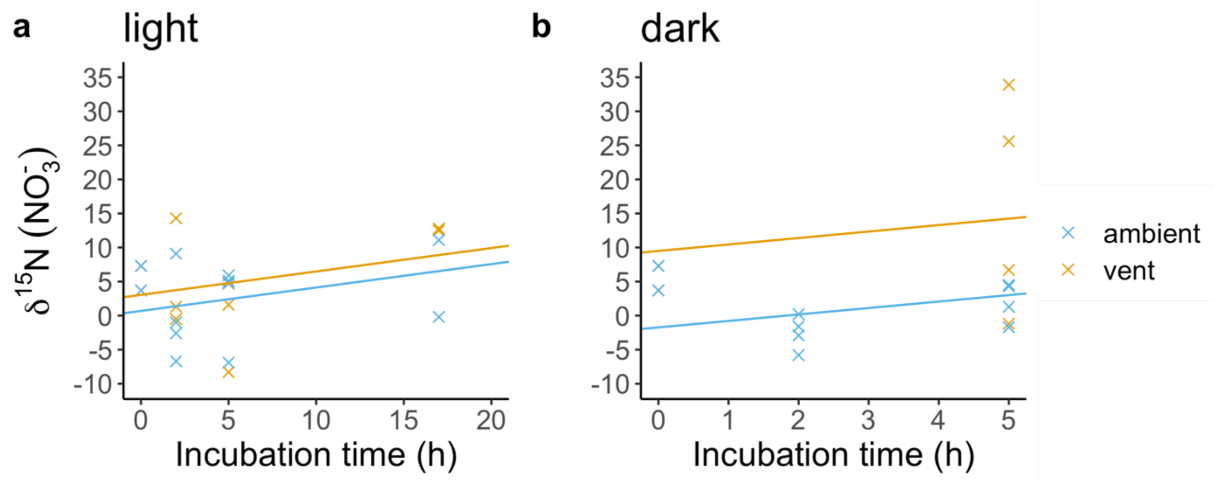


**Suppl. Figure 4.** ™$^{15}$N ($NO_3^-$) increase during light (a) and dark (b) incubations with seagrass leaf sections with epiphytes. Solid lines represent linear regressions.

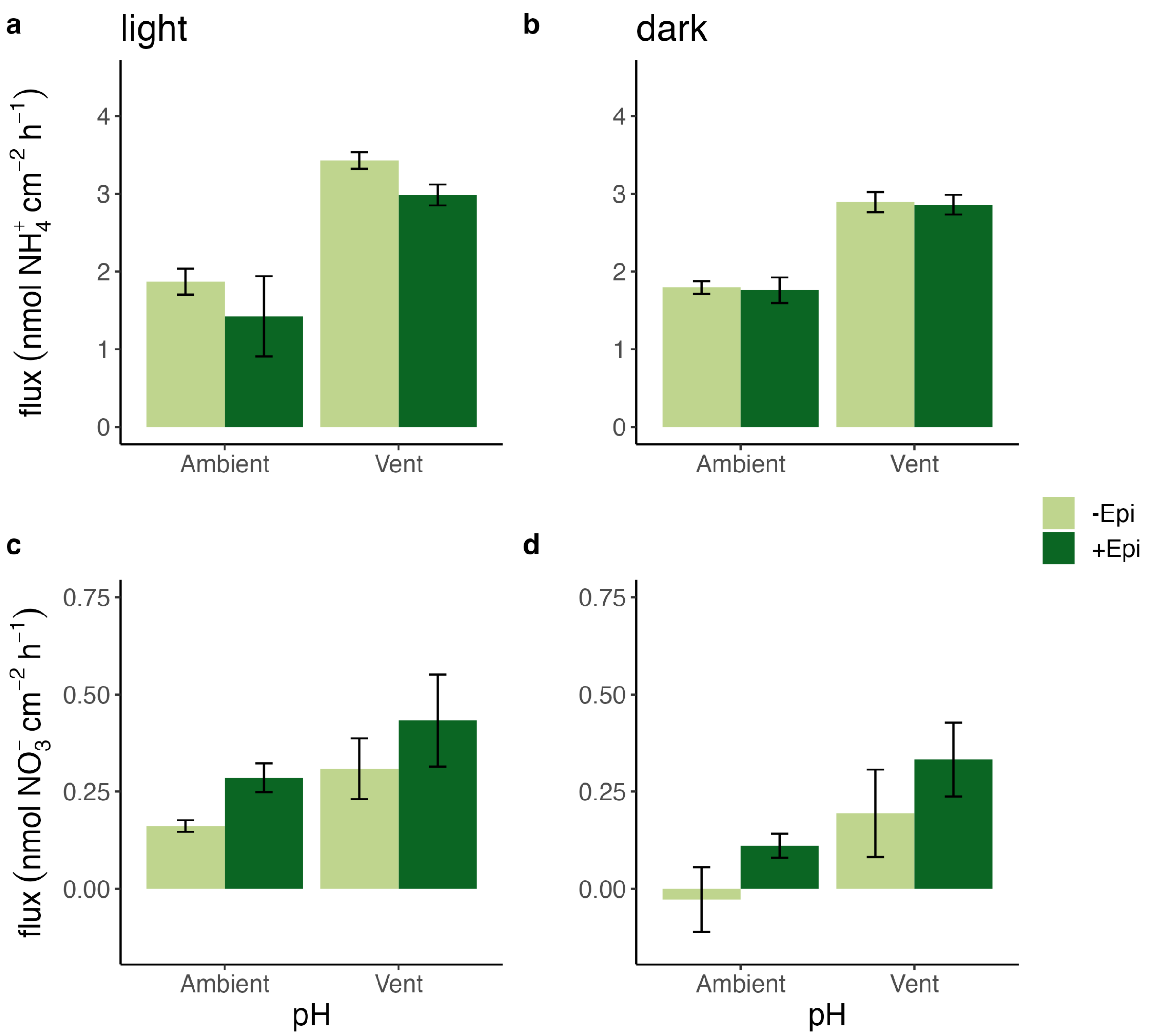


**Suppl. Figure 5.** Uptake of $NH_4^+$ (**a, b**) and $NO_3^-$ (**c, d**) during light (**a, c**) and dark (**b, d**) incubations with leaf sections from the ambient and vent site with (+Epi, n=4) and without epiphytes (-Epi, n=3). Error bars indicate mean ± SE.

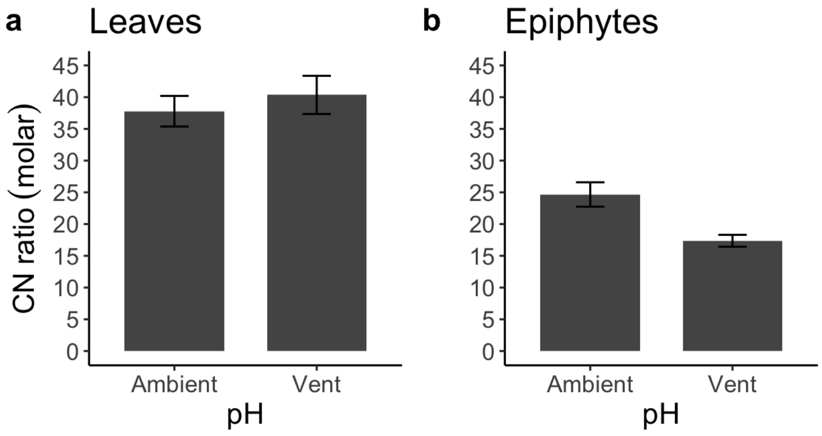


**Suppl. Figure 6.** C:N ratios of leaf sections (**a**) and epiphytes (**b**) from the ambient (*n* leaves = 14, *n* epiphytes =8) and vent site (*n* leaves = 14, *n* epiphytes =7). Since there were no differences between light and dark incubations, the samples were combined and treated as replicates. Error bars indicate mean ± SE.

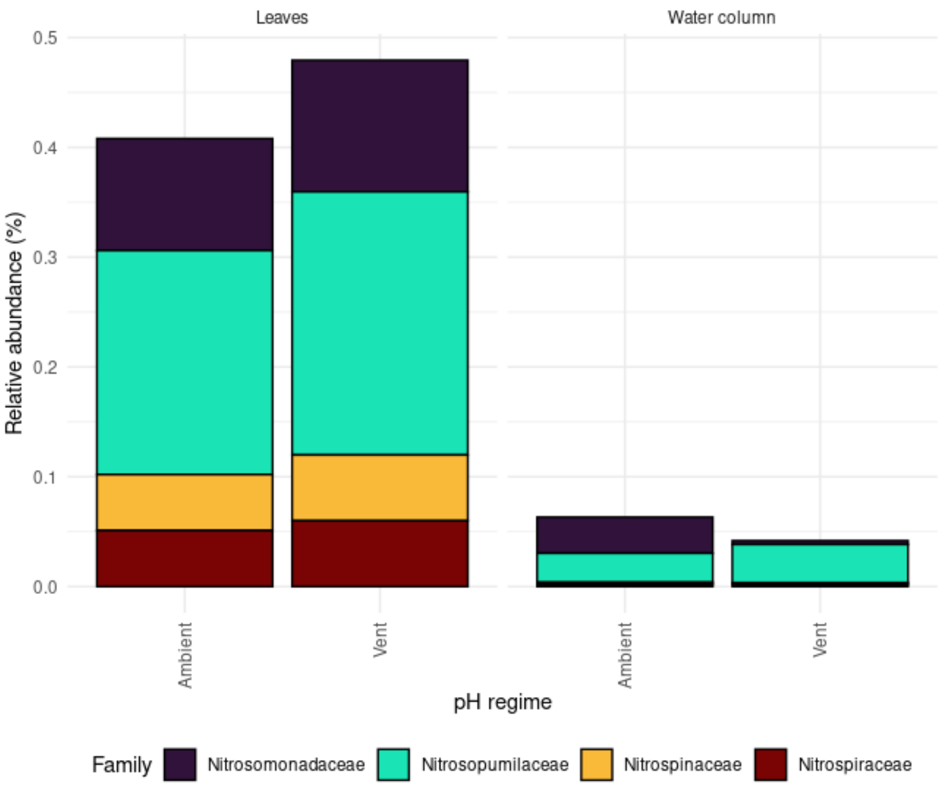


**Suppl. Figure 7.** Relative abundances of nitrifying prokaryotic taxa collapsed at the family level on leaves and water column samples from both pH regimes.

**Suppl. Table 1.** Environmental parameters (mean ± SE, n=3) measured at the vent and ambient pH site at Castello Aragonese.

| | Ambient pH | Vent pH |
|---|---|---|
| T (°C) | 23.94 ± 0.05 | 23.74 ± 0.01 |
| Light (Lux) | 10438 ± 872 | 16631 ± 628 |
| pH | 8.08 ± 0.04 | 7.80 ± 0.14 |
| DO (mg $L^{-1}$) | 9.15 ± 0.02 | 8.26 ± 0.02 |

Average temperature, light, and DO were continuously measured with data loggers during the sampling time between 11 am and 4 pm of the respective sampling day. PH was measured on 13.09.2019 with a pH logger (n ambient = 15, n vent = 8)

**Suppl. Table 2.** Permutation-based analysis of variance of the microbial communities associated with *P. oceanica* leaves, water column and pH regime.

| Source of variation | Degrees of freedom | Sum of squares | $R^2$ | Pseudo-F | P(>F) |
|---|---|---|---|---|---|
| pH regime | 1 | 821.5626 | 0.0641022 | 2.013119 | 0.1885 |
| Compartment | 1 | 8928.5753 | 0.6966495 | 21.878170 | 0.0010 |
| Treatment x Compartment | 1 | 617.6887 | 0.0481950 | 1.513556 | 0.2077 |
| Residual | 6 | 2448.6258 | 0.1910533 | NA | NA |
| Total | 9 | 12816.4525 | 1.0000000 | NA | NA |

**Suppl. Table 3.** Permutation-based analysis of variance of the nitrifying communities associated with *P. oceanica* leaves, water column and pH regime.

| Source of variation | Degrees of freedom | Sum of squares | $R^2$ | Pseudo-F | P(>F) |
|---|---|---|---|---|---|
| pH regime | 1 | 1.16E-08 | 0.0009 | 0.7777 | 0.9858 |
| Compartment | 1 | 1.44E-06 | 0.1125 | 9.6679 | 0.0001 |
| Treatment x Compartment | 1 | 2.25E-08 | 0.0017 | 0.1506 | 0.96 |
| Residual | 76 | 1.13E-05 | 0.8879 | NA | NA |
| Total | 79 | 1.28E-05 | 1.0000000 | NA | NA |

**Suppl. Table 3.** Morphological traits (mean ± SE) of *P. oceanica* from ambient and vent pH sites.

| | Ambient pH | Vent pH |
|---|---|---|
| Shoot density ($m^{-2}$) | 527.38 ± 110.90 | 1130.09 ± 234.24 |
| Leaf density ($m^{-2}$) | 4237.85 ± 515.01 | 7496.29 ± 674.21 |
| Leaf dry weight (g) | 0.041 ± 0.005 | 0.087 ± 0.013 |

**Suppl. Table 5.** $^{29}N_2$ and $^{29}N_2$ concentrations in the denitrification experiment at different incubation timepoints (mean ± SD).

| **Site** | **Timepoint** | **Incubation** | **Treatment** | **$^{29}N_2$ concentration (nmol/L)** | **$^{30}N_2$ concentration (nmol/L)** |
|---|---|---|---|---|---|
| Vent | T0 | | | 0.080 ± 0.015 | 0.263 ± 0.012 |
| Vent | T1 | light | +Epi | 0.075 ± 0.008 | 0.313 ± 0.024 |
| Vent | T1 | light | -Epi | 0.062 ± 0.006 | 0.355 ± 0.013 |
| Vent | T1 | dark | +Epi | 0.057 ± 0.008 | 0.323 ± 0.056 |
| Vent | T1 | dark | -Epi | 0.056 ± 0.008 | 0.249 ± 0.007 |
| Vent | T2 | light | control | 0.091 ± 0.018 | 0.019 ± 0.038 |
| Vent | T2 | dark | control | 0.042 ± 0.051 | -0.015 ± 0.054 |
| Vent | T2 | light | +Epi | 0.144 ± 0.036 | 0.369 ± 0.059 |
| Vent | T2 | light | -Epi | 0.105 ± 0.027 | 0.091 ± 0.021 |
| Vent | T2 | dark | +Epi | 0.097 ± 0.013 | 0.069 ± 0.008 |
| Vent | T2 | dark | -Epi | 0.098 ± 0.010 | 0.062 ± 0.012 |
| Ambient | T0 | | | 0.059 ± 0.015 | 0.302 ± 0.041 |
| Ambient | T1 | light | +Epi | 0.022 ± 0.080 | 0.231 ± 0.063 |
| Ambient | T1 | light | -Epi | 0.010 ± 0.114 | 0.229 ± 0.060 |
| Ambient | T1 | dark | +Epi | 0.025 ± 0.067 | 0.206 ± 0.062 |
| Ambient | T1 | dark | -Epi | -0.006 ± 0.096 | 0.209 ± 0.009 |
| Ambient | T2 | light | control | 0.104 ± 0.013 | 0.026 ± 0.006 |
| Ambient | T2 | dark | control | 0.097 ± 0.012 | 0.050 ± 0.023 |
| Ambient | T2 | light | +Epi | 0.122 ± 0.015 | 0.053 ± 0.037 |
| Ambient | T2 | light | -Epi | 0.072 ± 0.051 | 0.036 ± 0.043 |
| Ambient | T2 | dark | +Epi | 0.110 ± 0.014 | 0.024 ± 0.030 |
| Ambient | T2 | dark | -Epi | 0.035 ± 0.048 | -0.071 ± 0.104 |
